# The sympathetic nervous system enhances host immune responses to enteric bacterial pathogens

**DOI:** 10.1101/2025.03.20.642634

**Authors:** Emmy Tay, Michael Cremin, Kristina Sanchez, Ingrid Brust-Mascher, Colin Reardon

## Abstract

Mucosal immune responses to enteric bacterial infections are highly coordinated processes that orchestrate host protection while minimizing the potential for immune-triggered pathology. In the intestinal tract, bidirectional communication occurs between the nervous and immune systems to affect local immune responses by modulating the activity of resident and recruited immune cells, and indirectly on the supporting stromal cells. These neuroimmune signaling pathways that alter host defense have focused on specialized sensory innervation and the unique neurotransmitters released from them. Although the sympathetic nervous system has been established to induce a tissue-protective phenotype in subpopulations of neuron-associated macrophages in the small intestine, the role of these neurons during enteric bacterial infection was unknown. Using genetic labeling of activated neurons with *Arc*TRAP, we demonstrate that colonic infection induces activation of the rostral ventrolateral medulla, a major sympathetic center in the brainstem. The importance of peripheral sympathetic neurons was further demonstrated using chemical sympathectomy that significantly increased bacterial burden during *Citrobacter rodentium* (*C. rodentium*) infection. Increased bacterial burden was matched by a deficit in host protection due to reduced IFNγ production by colonic CD4+ T-cells. Sympathectomy, however, did not diminish the capacity to differentiate into IFNγ- or IL-17A- producing T-cells *in vitro*, suggesting that the lack of sympathetic innervation during infection may alter this process *in vivo* without causing sustained T-cell intrinsic defects. In assessing which receptors could mediate these effects, pharmacological antagonists selective for α- adrenergic receptors (αAR), but not β-adrenergic receptors, increased bacterial burden and reduced colonic IFNγ production. Using isolated cell types from the colon of uninfected and infected mice, we identified the αΑR subtypes expressed on immune and stromal cells, with significant upregulation of these receptors on T-cells during *C. rodentium* infection. Together these data demonstrate the unique role of the sympathetic nervous system and αAR in mucosal immune responses against enteric bacterial pathogens.

## Introduction

Protection from enteric bacterial pathogens requires a carefully choreographed immune response to provide protection while limiting the potential for collateral damage. Neurons and neuron-to-immune cell communication have now been well-established to control various aspects of immune function in a variety of organ systems^1,2^. This diverse array of neurons and neurotransmitters can control immune responses in the skin, lungs, small intestine, colon, spleen and lymph nodes during inflammation or infection^3–7^. In addition to the intrinsic enteric nervous system (ENS), the intestinal tract is also densely innervated by extrinsic neurons with cell bodies that reside in the sympathetic chain ganglia or dorsal root ganglia in the spinal cord. Although sensory neurons (marked by expression of the polymodal nociceptor, TRPV1) can release neuropeptides such as substance P to coordinate immune responses during enteric bacterial infection, the role of other types of innervation in modulating host immunity is only beginning to be addressed^4,5,8,9^. In the intestine, sympathetic innervation originates in the paravertebral chain ganglia, and projects to the myenteric plexus of the ENS and the lamina propria^10,11^. These neurons release neurotransmitters such as norepinephrine, vasoactive intestinal peptide (VIP), and co-release ATP to affect many physiological processes^11,12^.

Sympathetic neurons and the release of neurotransmitters have also been shown to alter host-microbe and host-pathogen interactions. The host response and the complex interplay of immune and non-immune cells during enteric bacterial infection *in vivo* have been robustly modeled by using the mouse-adapted pathogen *Citrobacter rodentium* (*C. rodentium*). This non-invasive bacterial pathogen transits along the gastrointestinal tract, infecting the cecum and colon by effacing the microvilli and attaching to the luminal surface of intestinal epithelial cells (IEC). This attachment permits the injection of bacterial effector proteins that allow for firm attachment and bacterial propagation by upregulating the type III secretion system. The early host response to this infection typically consists of activation of innate and resident immune cells including, macrophages, innate lymphoid cells (ILC), and recruitment of neutrophils and monocytes to the infected intestine. Type 3 ILCs (ILC3) produce high levels of IL-22 to increase expression of IEC antimicrobial peptides and are critical to host resistance and prevention of bacterial dissemination^13,14^. These early protective responses are superseded by the generation of CD4+ T-cells that are recruited and produce cytokines including IFNγ, IL-17A, and IL-22. These cytokines reinforce the ongoing inflammation and drive the expression of other host-protective factors, such as inducible nitric oxide synthase (Nos2) and MHCII, by a diverse array of cell types including IECs ^15–17^. Despite the requirement for T-cells in host protection, these proinflammatory responses can also enable bacterial survival due to the changing oxygenation of the colon, in addition to causing immunopathology^18^. Fine control of these inflammatory processes could allow for host protection from pathogens while limiting immune-induced damage.

The nervous system can modulate immune cell activation to either enhance or inhibit inflammation. Experimental activation of VIPergic neurons reduced the number of small intestinal and colonic ILC3. These mice were more susceptible to *C. rodentium* infection, due to direct inhibition of the ILC3 population^19^. In the small intestine, sympathetic neurons can also activate macrophages associated with the ENS to induce a reparative and neuronal protective state at baseline and following infection with *Salmonella typhimurium* (*S. typhimurium*) ^20^. The ability of sympathetic innervation to alter the trajectory of an immune response, and the specific cell types affected depend on the expression of α- and β-adrenergic receptors. In contrast to the β_2_-adrenergic receptors (β_2_AR) that have been described to reduce inflammation^21^, α-adrenergic receptors (αΑR) can increase immune cell activation and inflammation in a variety of settings^22,23^. Despite these observations, the role of sympathetic innervation and the αAR subtypes during enteric bacterial infection in the colon has not been described.

Here we demonstrate that infection with the enteric bacterial pathogen *C. rodentium* in mice induces neuronal activation of a major sympathetic control center in the brain. This sympathetic activity is host-protective and coordinates the immune responses during *C. rodentium* infection. Ablation of peripheral sympathetic neurons increased pathogen shedding, and bacterial burden, that correlated with decreased expression of many host-protective and proinflammatory cytokines. Significantly reduced colonic *Ifnγ* mRNA expression was matched with a reduced frequency of IFNγ+ colonic lamina propria CD4+ T-cells in sympathectomized infected mice compared to infected controls. These reduced responses were restricted to T-cells, as the recruitment of monocytes, macrophages, and neutrophils were unchanged in infected sympathectomized mice. To assess which receptors were required by sympathetic innervation to enhance host protection against *C. rodentium*, pharmacological antagonists of αAR and βAR were employed. In keeping with our data from chemical sympathectomy with 6OHDA, antagonism of αAR not only significantly increased bacterial burden, but significantly attenuated infection-induced IFNγ mRNA expression. Highlighting the unique role of αAR, these effects were not observed in either βAR antagonist-treated or β_2_AR KO mice. We further identified that T-cells not only express all the αAR subtypes, but that expression is significantly increased during *C. rodentium* infection. Together, these data highlight that sympathetic innervation and catecholamines can enhance host-protective response through αAR, but not βAR, during enteric bacterial infection. These findings further suggest that host immunity and inflammation could be fine-tuned using agonists or antagonists of these receptors to achieve clinically desired health goals.

## Methods & Materials

### Mice

Male and female C57BL/6J, C57BL/6N, B6.129(Cg)-Arctm1.1(cre/ERT2)Luo/J (Arc.CreERT2) and B6.Cg-Gt(ROSA)26Sortm75.1(CAG-tdTomato*)Hze/J (Ai9) mice were purchased from Jackson Laboratories (Bar Harbor, ME) to establish breeding colonies. *Arc*TRAP mice refer to the ArcCreERT2+/− Ai9+/− mice derived by crossing Arc.CreERT2 mice with Ai9 mice. Constitutive β_2_AR knockout (β_2_AR KO) mice on C57BL/6N background were purchased from the Mouse Biology Program (University of California, Davis). Male and female mice between 6 to 8 weeks old were utilized for all experiments. All procedures and protocols were approved by the Institutional Animal Care and Use at the University of California, Davis.

### Citrobacter rodentium infection

*C. rodentium* strain DBS100 was kindly provided by Dr. Andreas Baumler. Bacteria were grown on MacConkey agar plates at 37°C overnight. A single colony was selected for culture in Luria Broth (LB) at 37°C overnight, followed by sub-culturing at 37°C for 2 hours the next day. Mice were given LB or *C. rodentium* at a concentration of 10^8^ colony-forming units (CFU) through orogastric gavage. At 10 days post-infection (d.p.i.), fecal pellet and distal colonic tissue were collected and weighed to assess *C. rodentium* colonization. Samples were homogenized with a stainless-steel bead and a tissue lyser (Qiagen, Germantown MD), serially diluted, and plated onto MacConkey agar. After overnight incubation at 37°C, bacterial colonies were counted, and CFU was calculated per unit weight of each sample.

### Pharmacological agents

For neuronal labeling, cohorts of ArcTRAP mice received 4-hydroxytamoxifen (4-OHT) (Sigma, Germany) 10 days after LB or *C. rodentium* infection. A single dose of 4-OHT was given by intraperitoneal (i.p) injections at 50 mg/kg to label active neurons with tdTomato over 7 days, as previously described^24^. For chemical sympathectomy, cohorts of mice received saline or 6- hydroxydopamine (6OHDA) (Sigma) before *C. rodentium* infection. Three incremental doses of 6OHDA, from 100 mg/kg to 250 mg/kg, were given by i.p injection every 48 hours, followed by a 5-day rest period before *C. rodentium* infection. Additional cohorts of mice received saline, pan βAR antagonist (propranolol hydrochloride, 5mg/kg), or pan αAR antagonist (phentolamine mesylate, 10mg/kg) (Tocris, Minneapolis, MN) daily by intraperitoneal injection from the day of *C. rodentium* infection until the end of the experiment.

### Gastrointestinal motility

Distal colonic motility was measured in saline and 6OHDA-treated mice after 10 days of infection. This was performed by placing each mouse in a confined box and quantifying the number of fecal pellets within a 15-minute period.

### RNA isolation and Quantitative PCR

During necropsy, relevant tissues were collected, placed immediately into TRIzol (Invitrogen, Carlsbad, CA) and frozen at −80°C until RNA extraction was performed. Thawed tissues were homogenized with a stainless-steel bead and a tissue lyser (Qiagen). RNA extraction was performed as per manufacturer’s instructions (Invitrogen) and RNA concentration was quantified by nanodrop. iScript cDNA synthesis kit was used to synthesize cDNA from 1 μg of template RNA as per manufacture’s instruction (Bio-Rad, Hercules, CA). The resulting cDNA samples used to evaluate target gene expression by quantitative real-time PCR using PowerUp SYBR green reagent and primer pairs from Primerbank (**Table S1**) on QuantStudio6 Flex qPCR system (ThermoFisher Scientific, Waltham, MA).

**Table S1.**
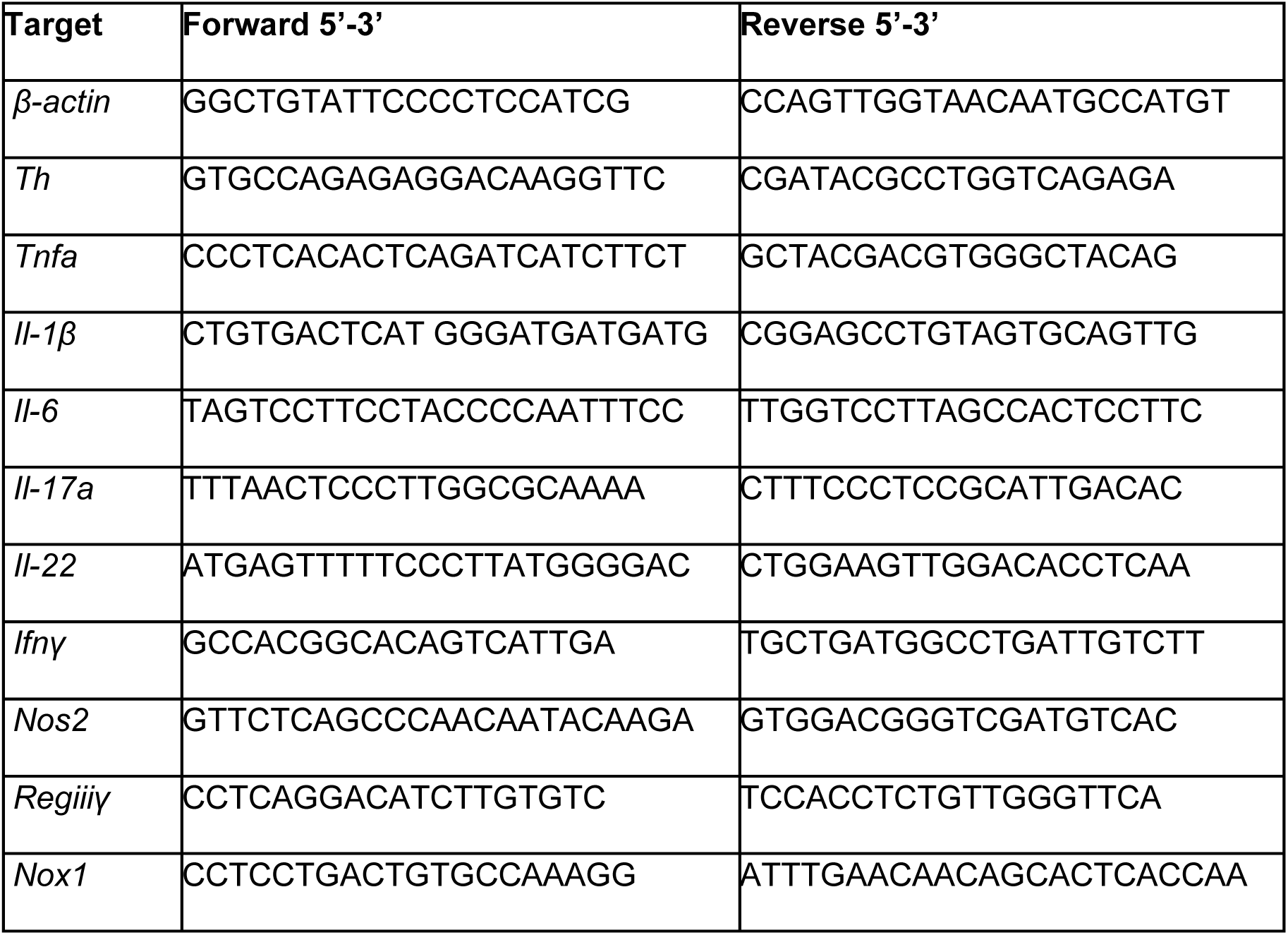
List of primers used for qPCR.

### Western blot

Colonic and splenic tissues were lysed with RIPA buffer (ThermoFisher Scientific) supplemented with protease inhibitor and phosphatase inhibitor cocktails (Roche, Indianapolis, IN). Tissue debris was removed by centrifugation at 15,000 xg for 15 minutes and protein concentrations were determined using the DCTM protein assay kit (Bio-Rad). Equal amount of protein lysates were loaded into each well and the samples were resolved by SDS-PAGE (Bio- Rad), followed by transfer onto nitrocellulose membrane (Bio-Rad) for incubation with anti- Tyrosine Hydroxylase (Cat #AB152, Sigma) and anti-Actin (Cat # 3700S, Cell Signaling Technology, Denvers, MA), followed by HRP-linked Goat anti-Rabbit (Cat # 7074P2, Cell Signaling Technology) and HRP-linked Goat anti-Mouse (Cat # 7076P2, Cell Signaling Technology). The blots were developed using the SuperSignal West Dura Substrate (ThermoFisher Scientific) and acquired by the ChemiDoc imaging system (Bio-Rad).

### Histology

During necropsy, relevant tissues were collected, fixed in 10% buffered formalin, processed and embedded in paraffin according to standard protocols. Paraffin embedded samples were sectioned into 6 μm sections with a microtome for histopathology or confocal microscopy. Sections for histopathology were deparaffinized, rehydrated and stained with hematoxylin and eosin following standard protocols. Colonic epithelial cell hyperplasia was assessed by bright field microscopy to allow for measurements of crypt length in at least 20 well-orientated crypts per sample using ImageJ software (NIH).

### Cryosection

*Arc*TRAP mice were perfused, brains collected and post-fixed in 4% PFA for 24 h, followed by incubation in 30% sucrose in PBS for another 48 h. Brains were subsequently embedded in OCT medium (ThermoFisher Scientific) and stored at −80°C overnight. Coronal sections (25 μm) were cut using a cryostat (Leica, Germany). Sections of NTS were collected around Bregma −7.48mm while sections of RVLM were collected around Bregma −6.84 mm according to the Mouse Brain Atlas.

### Confocal microscopy

Slides with paraffin tissue sections were prepared as described above, followed by deparaffinization and rehydration, and antigen retrieval was performed in citrate buffer (10 mM, pH 6.0) for 30 minutes at 95°C. Sections were then incubated in blocking buffer (5% BSA with normal donkey and goat serum) for 1 h at RT, followed by incubation in anti-Ki67 (Cat #LS- C141898, LS Bio, Newark, CA) overnight at 4°C. Sections were washed extensively with washing buffer (0.01% Tween-20 in TBS), followed by incubation in Donkey anti-Rabbit Alexa Fluor 546 (Cat #10040, Invitrogen) for 1h at RT. After another round of extensive washing, sections were counterstained with nuclear dye DAPI and mounted in Prolong Gold antifade reagent (Invitrogen). For sections stained with anti-CDH1 (Cat # CP1921, ECM Biosciences, Aurora, CO), a mouse-on-mouse kit was used in addition to the procedure described above as per the manufacturer’s instructions (Vector laboratories, Burlingame, CA). Stained slides were imaged on a Leica SP8 STED 3X confocal microscope with a 20X/0.75 NA or 40X/1.3 NA objectives. Whole tissue sections were acquired using a tiling approach, with 10% overlap between adjacent images. Files were processed by Imaris Stitcher (Bitplane Scientific) to recreate a composite image of the sample. DAPI+ CDH1+ Ki67+ cells were determined and quantified using Imaris (Oxford Instruments, UK), and divided by the number of crypts defined within the region^9^.

### T-cell enrichment

Mesenteric lymph nodes were sterilely excised during necropsy, placed on a 100 μm filter and dissociated using the plunger of a syringe. Single-cell suspensions were obtained by washing extensively with stain buffer (2% FBS in PBS), followed by treatment with Ack lysis solution for 5 minutes at RT. These cells were then incubated in Fc block (anti-CD16/32, 10 µg/ml, Tonbo Biosciences, San Diego, CA) for 15 minutes on ice. Antibodies for cell surface staining (**Table S2**) were added for another 30 minutes on ice, followed by washing in stain buffer. The following biotinylated antibodies were added at 1:50 dilution: anti-CD161 (Clone # PK136, Cat # 30- 5941), anti-CD11c (Clone # N418, Cat # 30-0114), anti-Ly6G (Clone # RB6-8C5, Cat # 30- 5931), anti-TER119 (Clone # TER-119, Cat # 30-5921), anti-CD11b (Clone # M1/70, Cat # 30-0112), anti-CD8 (Clone # 53-6.7, Cat # 30-0081) and anti-B220 (Clone # RA3-6B2, Cat # 30- 0452) from Tonbo Biosciences. Cells were then washed and incubated in magnetic streptavidin beads (BD Biosciences, Franklin Lakes NJ) according to the manufacturer’s protocol. Cells were placed in BD IMAG Cell Separation Magnet (BD Biosciences) for 8 minutes and the negative fraction was taken and placed into a fresh tube on the IMAG. This process was repeated for two additional magnetic enrichment steps. T-cell purity was confirmed >90% by flow cytometry.

### *In vitro* T-cell differentiation and ELISA

Negatively enriched CD4+ T-cells were plated on a round bottom 96-well plate pre-treated with 2 µg/mL anti-CD3 (Cat # 40-0032, Tonbo Biosciences) for 2 h 37°C. Cells were cultured with or without 0.5 µg/mL anti-CD28 (Ca t# 40-0281, Tonbo Biosciences) and in Th1 or Th17 skewing culture conditions. Th1 differentiation media consisted of 1 µg/mL anti-murine IL-4 (Cat # 16- 7041, Invitrogen), 5 ng/mL murine IL-2 (Cat # 212-12, Peprotech Cranbury, NJ), and 10 ng/mL murine IL-12p40 (Cat # 210-12, Peprotech). Th17 differentiation media consisted of 1 µg/mL anti-murine IFNγ (Cat # 16-7312, Invitrogen), anti-murine IL-4 (Cat # 16-7041, Invitrogen), 1 µg/mL anti-murine IL-2 (Cat # 14-7022, Invitrogen), 20 ng/mL murine IL-6 (Cat # 216-16, Peprotech), and 1 ng/mL murine TGF-β1 (Cat # 7666-MB, R&D Systems Minneapolis, MN). Cells were incubated at 37°C for 96 h, then stimulated with 1X Cell Stimulation Cocktail (eBioscience, San Diego CA) for 4 h. Supernatant was collected for ELISA. Untreated 96 flat bottom plates were coated with capture antibody for IFNγ (Cat # 88-7314-22, Invitrogen) or IL- 17A (Cat # 88-7371-88, Invitrogen) as per manufacturer’s protocols. Plates were read on a BioTek Synergy HTX Multi-mode Plate Reader using the Gen5 application. The standard curve was determined using 4-paramter logistic fit.

### Isolation of cells from the colon

Colonic tissues were collected during necropsy and reduced into single-cell suspensions using the lamina propria dissociation kit (Miltenyi Biotec, Gaithersburg, MD) with a gentleMACS tissue dissociator as per manufacturer’s protocol. In brief, colonic tissues were opened longitudinally and cut into 1 cm long pieces. Intestinal epithelial cells (IEC) were removed by gentle agitation in HBSS supplemented with 10 mM HEPES, 5 mM EDTA and 5% FBS, washed and stored in TRIzol for RNA extraction. Colonic tissues were then digested using MACS digestion enzyme mix and reduced to a single-cell suspension by the gentleMACS tissue dissociator. The resulting cell suspensions were passed through a 100 µm strainer and before proceeding with staining for flow cytometry.

### Flow cytometry

Single-cell suspensions were manually counted with a hemocytometer and incubated in Fc block (anti-CD16/32, 10 µg/ml, Tonbo Biosciences) for 25 minutes on ice, followed by incubation in antibody cocktail for surface staining (**Table S2**) for 30 minutes on ice. Cell viability was determined using fixable live/dead aqua as per manufacturer’s instructions (Cat # L34957, ThermoFisher Scientific). After incubation with an antibody cocktail, cells were fixed with BD Cytofix (BD Biosciences) for 25 minutes on ice. All flow cytometry data was acquired on a LSRII (BD Biosciences) using DIVA software and analysed using the FlowJo software (Becton Dickinson, Eugene, OR).

### Intracellular staining

Single-cell suspensions were manually counted with a hemocytometer and incubated in RPMI media with 10% FBS, 1% Penicillin/ Streptomycin, 2 mM L-glutamine, BD GolgiPlug (Cat # 555029, BD Biosciences), and stimulated with 1X Cell Stimulation Cocktail (eBioscience) at 37°C for 4 h. After extensive washing with stain buffer (2% FBS in PBS), surface staining was performed as per the procedures described above, followed by incubation in BD Cytofix/ Cytoperm Fixation/ Permeabilization Solution kit (Cat # 554714, BD Biosciences) for 1 h at RT. Intracellular staining was performed by incubating single cells in antibodies for cytokines (**Table S2**) diluted in in 1X BD Perm/ Wash Buffer (Cat # 554723, BD Biosciences) for 1 hour at RT. For FoxP3 staining, cells were fixed and permeabilized using FoxP3 Transcription Factor Staining Kit (Cat # 00-5523, ThermoFisher Scientific) for 1 h at 4°C, followed by staining with anti-FoxP3 for 1 h at RT.

**Table S2.** Antibodies used for flow cytometry.

| Antibody | Conjugate | Manufacture | Clone | Catalog No. |
| --- | --- | --- | --- | --- |
| CD45 | eFluor450 | Invitrogen | 30-F11 | 48-0451-82 |
| CD3 | APC | Tonbo Biosciences | 145-2C11 | 20-0031 |
| CD4 | FITC | BD Biosciences | RM4-5 | 553047 |
| B220 | PerCP Cy5 | BioLegend | RA3-6B2 | 103210 |
| Ly6G | FITC | Tonbo Biosciences | 1A8 | 35-1276-U100 |
| Ly6C | PE Cy7 | Invitrogen | HK1.4 | 25-5932-82 |
| CD64 | PE | Miltenyi Biotec | REA286 | 130-118-684 |
| CD11b | Alexa Fluor 700 | Invitrogen | M1/70 | 45-0112 |
| CD31 | APC Cy7 | BioLegend | MEC13.3 | 102533 |
| gp38 | Alexa Fluor 488 | BioLegend | 8.1.1 | 127405 |
| IFNγ | PE Cy7 | Invitrogen | XMG1.2 | 25-7311-82 |
| IL-17A | PE | BD Biosciences | TC11-18H10 | 561020 |
| IL-22 | PerCP eFluor 710 | Invitrogen | 1HBPWSR | 46-7221-82 |
| FoxP3 | PE Cy7 | Invitrogen | FJK-16s | 25-5773 |

### mRNA expression determination in FACS-sorted cells

LB and *C. rodentium*-infected mice were euthanized at 10 d.p.i. Colons were collected and processed into single-cell suspensions, as described above. Isolated single cells were stained with antibodies **(Table S2)**: anti-CD45, anti-CD3, anti-Ly6G, anti-gp38, anti-CD31, and Live/ Dead Fixable Aqua. Blood endothelial (BEC: CD45-, CD31+, gp38-), lymphatic endothelial (LEC: CD45-, CD31+, gp38+), neutrophils (CD45+, CD3-, Ly6G+) and T-cells (CD45+, CD3+, Ly6G-) were sorted into separate tubes using an Astrios Cell Sorter (Beckman Coulter Brea, CA). RNA was extracted and processed for qPCR analysis using the CellAmp Direct TB Green RT-qPCR Kit (Takara Bio, San Jose, CA, Cat# 3735A) according to the manufacturer’s protocol.

### Statistics

Statistical analysis of all data was performed using Prism 10.0 (GraphPad, La Jolla, CA) with a Student’s t test or one-way ANOVA followed by post-hoc analysis with Tukey’s multiple comparison test. Individual data points are presented as mean ± standard deviation of the mean.

## Results

Enteric bacterial infection with *C. rodentium* activates sympathetic brain center and is worsened by the ablation of sympathetic innervation.

The ability of enteric bacterial infection with *C. rodentium* to evoke neuronal activation in discrete brain regions was assessed using the *Arc*TRAP method to genetically mark neurons with temporal control^24^. Administration of 4-hydroxytamoxifen (4-OHT) to Arc.CreERT2+ LSL-TdTomato mice 10 days post-infection (d.p.i.) (**Fig 1A**) increased tdTomato expression in discrete loci within the brain stem compared to non-infected controls. These regions included the nucleus tractus solitarius (NTS), a major sensory center in the brainstem (**Fig S1A**), as well as the rostral ventrolateral medulla (RVLM), a major sympathetic control center (**Fig 1B**).

**Figure 1.**
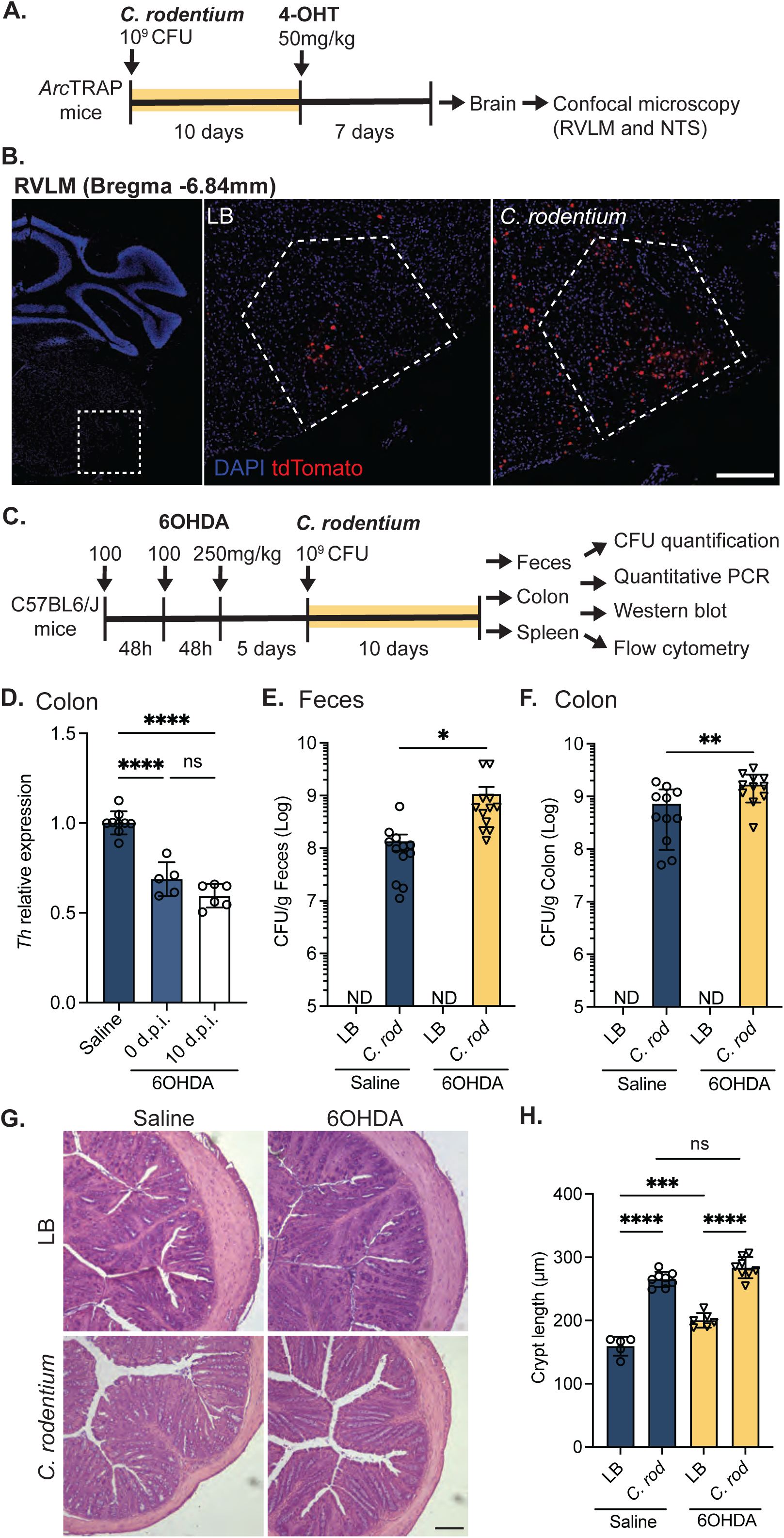
*C. rodentium* infection activates neurons within sympathetic brain regions, and is worsened by the ablation of sympathetic innervation. The experimental timeline detailing how Arc.CreERT2 LSL-tdTomato (*Arc*TRAP) mice were infected with *C. rodentium* (10^9^ CFU, orogastric gavage), followed by 4-hydroxytamoxifen (4-OHT, 50 mg/kg, i.p.) administered at 10 days post-infection (d.p.i.) to label active neurons with tdTomato **(A)**. Infection with *C. rodentium* increased the number of activated neurons (DAPI+ tdTomato+) within the rostral ventrolateral medulla (RVLM) compared to non-infected control mice **(B)**. Peripheral sympathectomy was performed according to the experimental timeline **(C)** with depletion confirmed by assessing colonic Tyrosine Hydroxylase (*Th*) mRNA expression **(D)**. Loss of peripheral sympathetic innervation significantly increased *C. rodentium* (“C rod”) bacterial burden at 10 d.p.i by quantification of colony-forming units (CFU) in feces **(E)** and colon **(F)**. Infection-induced colonic crypt hyperplasia was assessed from H&E images **(G)** by quantification of crypt length 10 d.p.i. in intact or sympathectomized uninfected and infected mice **(H)**. Results are from individual mice, mean ± SD. * P≤0.05, ** P ≤0.01, **** P ≤0.0001. One-way ANOVA with Tukey’s post-hoc test. Scale bar = 100 µm.

To determine the role of peripheral sympathetic innervation during *C. rodentium* infection, chemical sympathectomy with 6-hydroxydopamine (6OHDA) was performed prior to the infection as illustrated in **Fig 1C**. Ablation of sympathetic innervation was confirmed by reduced Tyrosine Hydroxylase (TH) mRNA and protein in the colon (**Fig 1D and S1B**) and the spleen (**Fig S1C&D**) throughout the time course of our experimental infection regimen. Mice that were sympathectomized and infected exhibited significantly increased fecal, and colon-associated *C. rodentium* burden 10 d.p.i. compared to infected mice with an intact sympathetic nervous system (**Fig 1E&F**). These increases in bacterial burden were not associated with increased crypt hyperplasia (**Fig 1G&H**), a prominent pathological feature of *C. rodentium* infection. Similarly, IEC proliferation was increased by *C. rodentium* infection but was not significantly different between control and sympathectomized mice **(Fig S1E&F)**. We further assessed colonic motility as a possible mechanism for increased bacterial burden but observed no differences between the control and sympathectomized mice (**Fig S1G&H**), indicating that increased *C. rodentium* was not simply due to reduced intestinal motility after sympathectomy. Taken together, these data demonstrate that sympathetic innervation could play a critical role in mediating host responses during an enteric infection with *C. rodentium*.

### Loss of sympathetic innervation reduces select aspects of host-protective immunity

With increased bacterial burden, we assessed if the loss of sympathetic innervation altered the immune response during enteric *C. rodentium* infection. Mice that were sympathectomized and infected had significantly reduced colonic *Ifnγ* and *Il17a*, but not *Il22* mRNA expression compared to non-sympathectomized infected controls (**Fig 2A-C**). Infection-induced expression of *Tnfα* and *Il1β* mRNA were also significantly reduced with sympathectomy compared to infected controls **(Fig 2D&E)**, but no significant differences were observed with *Il6* expression (**Fig 2F**). In assessing known host-protective genes, sympathectomy significantly reduced mRNA expression of the IFNγ regulated genes *Nos2* and *Nox1*, (**Fig 2G&H**), without affecting the expression of the antimicrobial gene *Reg3g* (**Fig 2I**). These data demonstrate that sympathetic innervation enhances specific aspects of host-protective responses induced during enteric infection with *C. rodentium*.

**Figure 2.**
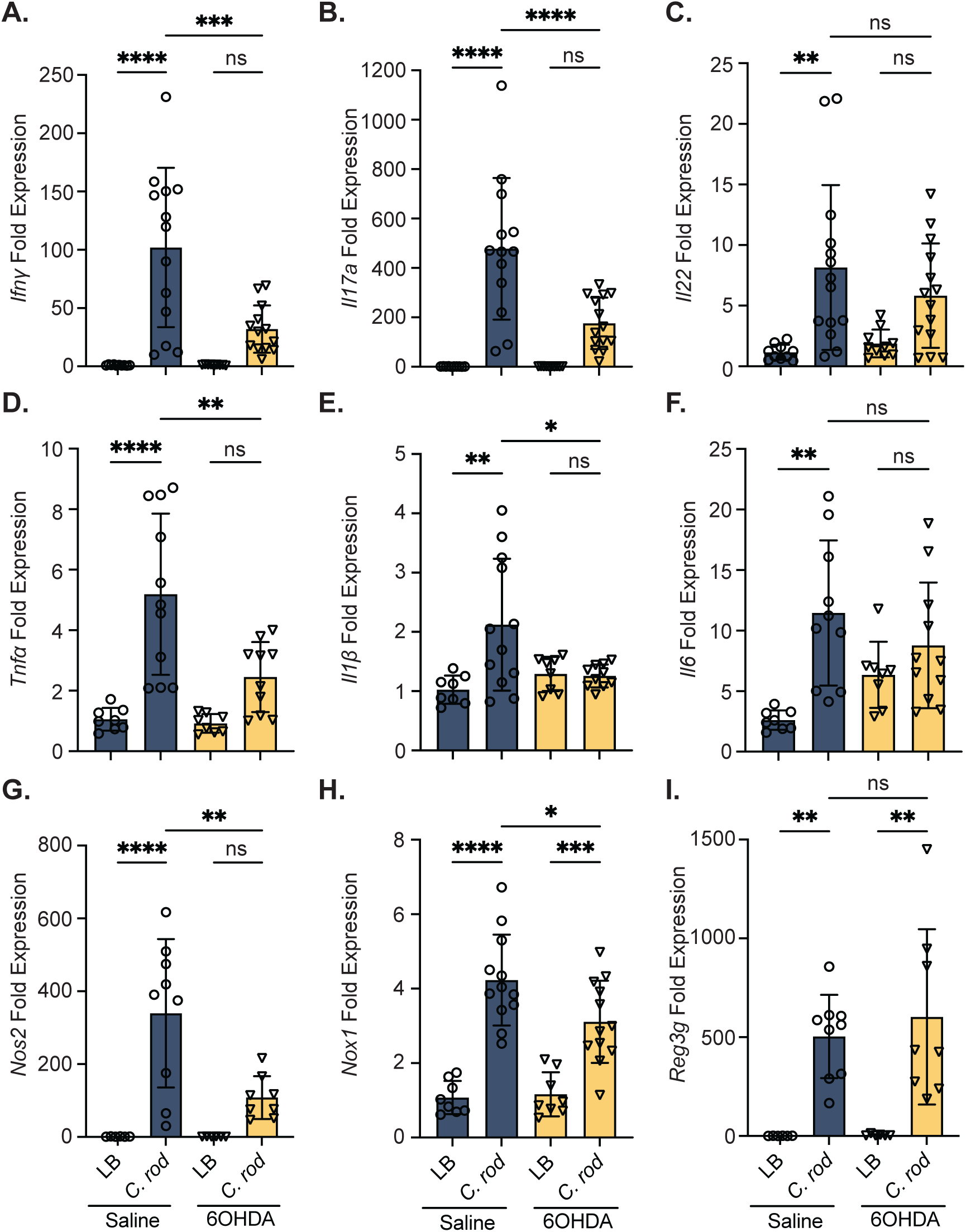
Loss of sympathetic innervation reduces host protective immune responses during *C. rodentium* infection. Expression of the host protective cytokines *Ifnγ* **(A)**, *Il17a* **(B)**, and *Il22* **(C)** in the colon of intact or sympathectomized, uninfected and *C. rodentium* (10 d.p.i.) infected mice were assessed by RT-qPCR. Colonic expression of the inflammatory cytokines *Tnfα* **(D)**, *Il1β* **(E)**, *Il6* **(F)**, and protective factors *Nos2* **(G)**, *Nox1* **(H)**, and *Reg3g* **(I)** were assessed from these intact sympathectomized uninfected and *C. rodentium* infected mice (10 d.p.i.). Results are from individual mice, mean ± SD. ns = not significant, * P≤0.05, ** P ≤0.01, *** P ≤0.001, **** P ≤0.0001. One-way ANOVA with Tukey’s post-hoc test.

### Sympathectomy reduces CD4+ T-cell responses during *C. rodentium* infection

As peripheral sympathectomy reduced mRNA expression of cytokines and antimicrobial genes critical for host protection, we assessed if colonic recruitment of various immune cell populations was reduced using flow cytometry (**Fig S2A**). As expected, infection with *C. rodentium* significantly increased the frequency and the number of neutrophils and monocytes in the colon, however, this was not different between the intact and sympathectomized mice **(Fig S2B-G)**. In assessing lamina propria lymphocytes (**Fig S3A**), the frequency and absolute numbers of T-cells in the colon were significantly increased by *C. rodentium* infection as expected, and sympathectomy with 6OHDA further increased colonic T-cell recruitment 10 d.p.i. **(Fig 3A&B)**. Intracellular staining further revealed no significant difference in the frequency of colonic Foxp3+ T-cells in sympathectomized versus intact mice irrespective of infection **(Fig 3C)**. When assessing key cytokines elicited by *C. rodentium*, we found significantly reduced CD4+ T-cells producing IFNγ in infected sympathectomized compared to infected controls mice **(Fig 3D&S3B)**, but no difference in IL-17A or IL-22 producing CD4+ T-cells between the two groups **(Fig 3E&F, S3C&D)**.

**Figure 3.**
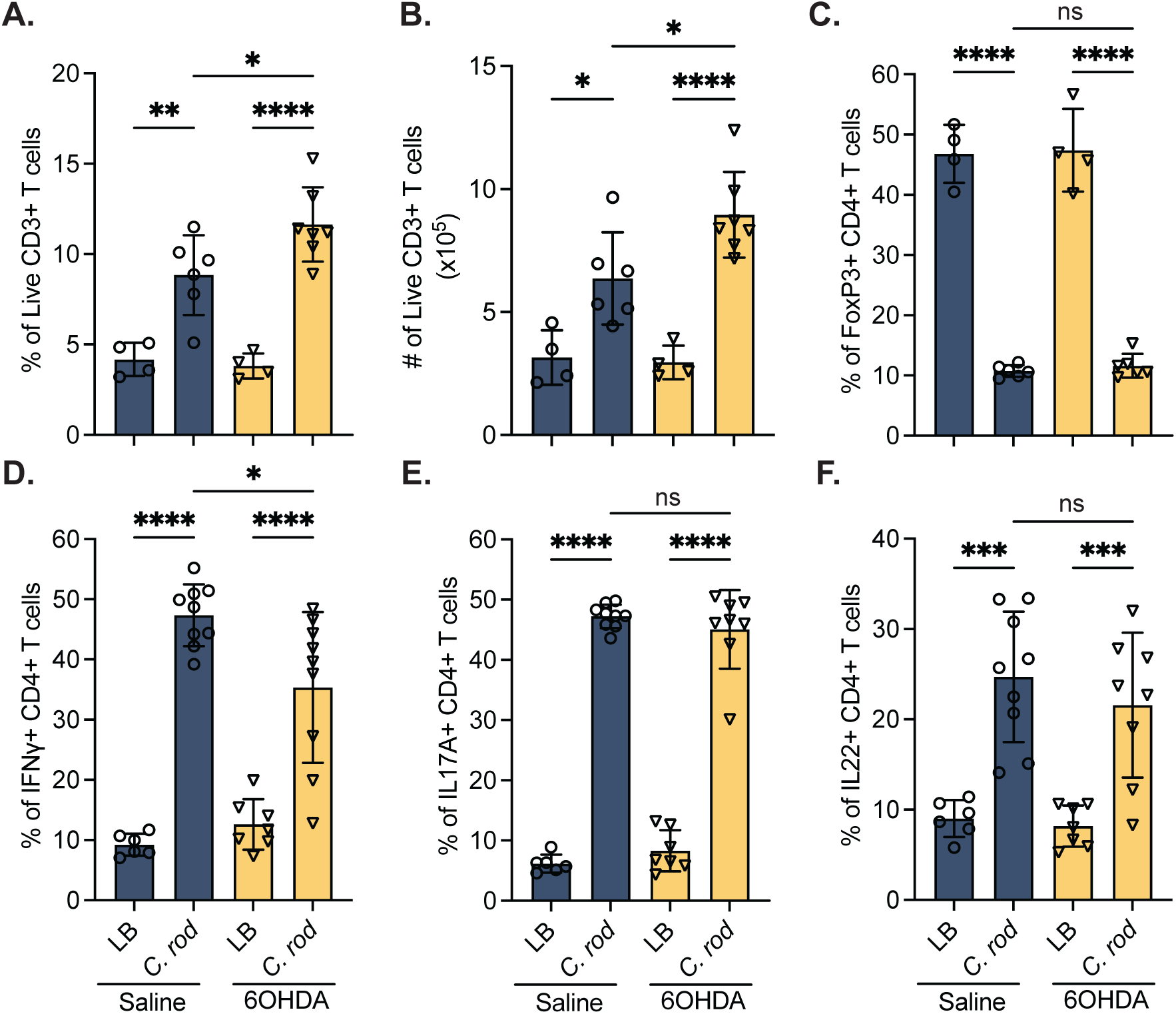
Sympathectomy reduces CD4+ T-cell responses during *C. rodentium* infection. Colonic recruitment of T-cells in control and sympathectomized uninfected or *C. rodentium*-infected mice was determined by flow cytometry. Frequency **(A)** and absolute numbers **(B)** of live CD3+ T-cells in the colonic lamina propria of control or 6OHDA-treated mice at 10 days post-infection. Intracellular cytokine staining for CD3+ CD4+ T-cells that express FoxP3 **(C)**, IFNγ **(D)**, IL-17A **(E)** and IL-22 **(F)** were quantified by flow cytometry. Results are from individual mice, mean ± SD. ns = not significant, * P≤0.05, ** P ≤ 0.01, *** P ≤0.001, **** P ≤0.0001. One-way ANOVA with Tukey’s post-hoc test.

To determine if sympathectomy prevented T-cells from committing to a Th1 effector phenotype, we performed *in vitro* differentiation assays using T-cells harvested from the mesenteric lymph node (MLN) of intact and sympathectomized mice. Differentiation to Th1 (IFNγ-secreting), or Th17 (IL17A-secreting) was not significantly different in T-cells originating from the MLN of intact versus sympathectomized mice **(Fig S3E&F)**. Together, these data demonstrate that sympathetic innervation contributes to specific aspects of the host immune response through enhancing IFNγ production by colonic CD4+ T-cells, but that lack of innervation does not permanently impinge on the ability of T-cells to differentiate into Th1 or Th17 T-cells.

### Development of host-protective T-cell responses is aided by α-adrenergic receptor signaling

To decipher the catecholamine receptors that aid host-protective responses during *C. rodentium* infection, a pharmacological antagonist approach was used. The role of αAR was determined by the administration of the non-subtype selective αAR antagonist, phentolamine mesylate, daily over the 10-day infection period **(Fig 4A)**. Treatment with αAR antagonist significantly increased the bacterial burden in the feces and colon **(Fig 4B&C)**, and significantly reduced *Ifnγ* mRNA expression **(Fig 4D)** at 10 d.p.i. compared to vehicle-treated infected mice. However, no significant difference in *Il17a* mRNA expression was observed between the infected groups **(Fig 4E)**. To determine which cell types receive sympathetic input through αAR, mRNA expression of these αAR subtypes were assessed in FACS-sorted T-cells (live, CD45+, CD3+) and neutrophils (live, CD45+, Ly6G+) from the colon of control and *C. rodentium* 10 d.p.i. Expression of *Arda1a*, *Arda1b*, *Arda1d*, *Arda2a*, *Arda2b* and *Arda2c*, were detected in control mice, and were significantly upregulated in colonic T-cells of *C. rodentium* infected mice at 10 d.p.i. (**Fig 4F-K**). Although these receptors were detected in colonic neutrophils, expression was not significantly altered by *C. rodentium* infection **(Fig 4L-Q)**. Expression of αAR subtypes was not restricted to immune cells, as we observed baseline expression in blood endothelial cells, with significant increases in *Adra1b, Adra2a, Adra2b, Adra2c* with *C. rodentium* infection at 10 d.p.i. (**Fig S5B,F,J,N,R,V**). Expression of these αAR receptors was also found in other stromal (**Fig S5A,E,I,M,Q,U**), and lymphatic endothelial cells (**Fig S5C,G,K,O,S,W)**, but was not significantly altered by *C. rodentium* infection. Despite significant decreases in αAR mRNA in intestinal epithelial cells 10 d.p.i. with *C. rodentium*, all of these receptor subtypes were minimally expressed at baseline (**Fig S5D,H,L,P,T,X**).

**Figure 4.**
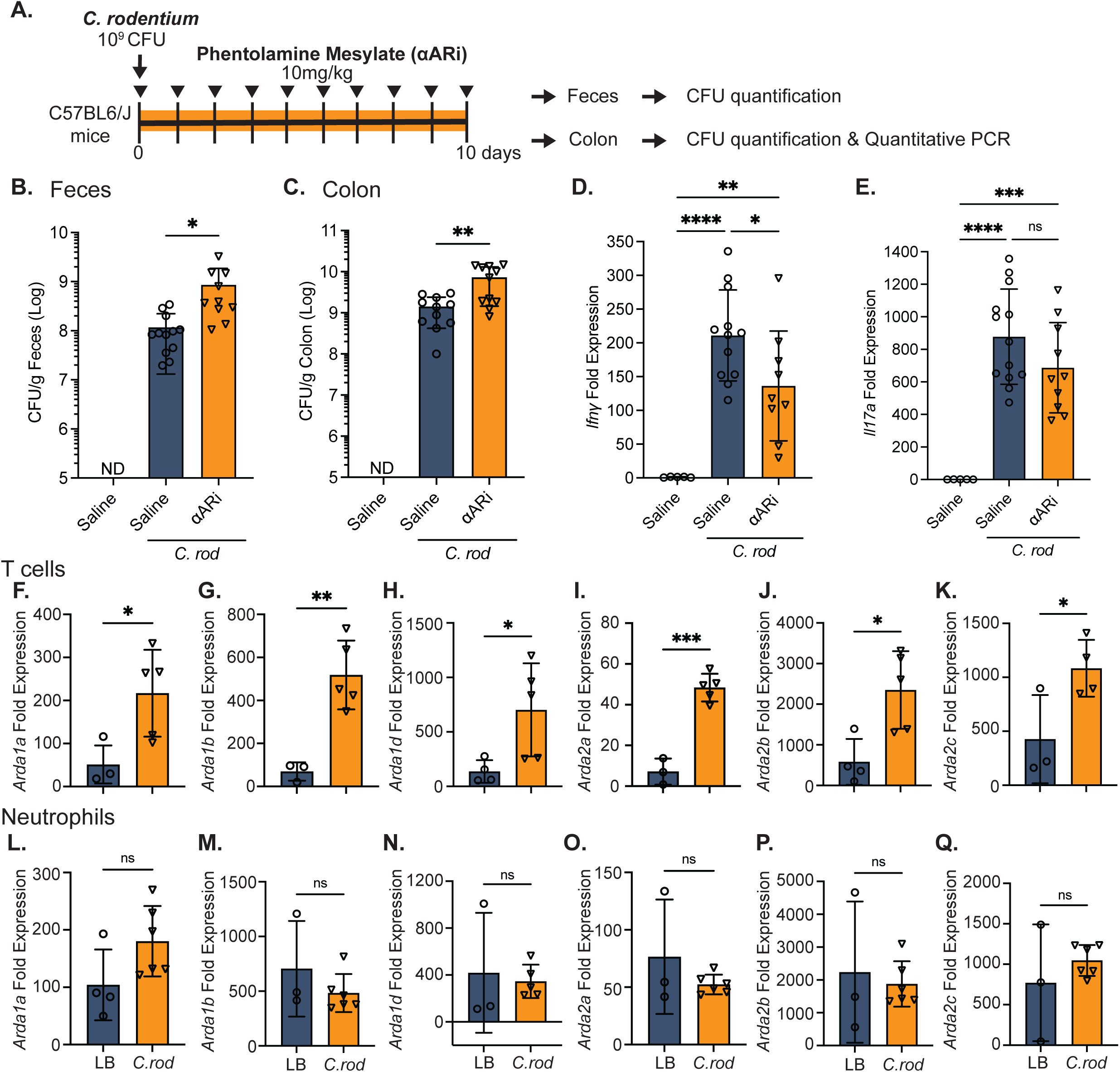
α-adrenergic receptor signaling aids the development of host-protective T-cell responses. To determine the role of α-adrenergic receptor (αAR) signaling during *C. rodentium* infection, mice received vehicle or the non-selective αAR antagonist (αARi), Phentolamine Mesylate (10 mg/kg i.p., one dose daily) after *C. rodentium* infection, with tissues collected 10 days post-infection (d.p.i.) **(A)**. Bacterial burden was assessed in vehicle and αARi uninfected and infected mice by quantification of *C. rodentium* in feces **(B)** and colon **(C)**. The colonic mRNA expression of *Ifnγ* **(D)** and *Il17a* **(E)** in uninfected and infected saline or αARi-treated mice at 10 d.p.i. was assessed by RT-qPCR. FACS sorted colonic T-cells (live, CD45+, CD3+) and neutrophils (live, CD45+, Ly6G+) were assessed for expression of αAR subtypes encoded by *Adra1a* **(F&L)**, *Adra1b* **(G&M)**, *Adra1d* **(H&N)**, *Adra2a* **(I&O)**, *Adra2b* **(J&P)** and *Adra2c* **(K&Q)** from LB and C. rodentium infected mice at 10 d.p.i. Results are from individual mice, mean ± SD. ns = not significant, * P≤0.05, ** P ≤0.01, **** P ≤0.0001. One-way ANOVA with Tukey’s post-hoc test.

To assess the role of β-adrenergic receptors (βAR), the non-subtype selective βAR antagonist, propranolol hydrochloride was administered daily over the 10-day infection period **(Fig S5A)**. Pharmacological antagonism of βAR significantly increased bacterial burden in feces but not in the colon **(Fig S5B&C)**. In addition, there was no change in *Ifnγ* or *Il17a* mRNA expression compared to vehicle-treated infected controls **(Fig S5D&E)**. Similar findings were observed in β_2_-adrenergic receptor knockout (β_2_AR KO) mice that showed increased bacterial burden **(Fig S5G&H)**, but no significant differences in the expression of *Ifnγ* or *Il17a* between genotypes during infection **(Fig S5I&J)**. These data demonstrate that host-protective immune responses, including the production of IFNγ during enteric bacterial infection, is enhanced through α-adrenergic, but not β-adrenergic signaling.

## Discussion

As a major interface with the external environment, the intestine is continuously exposed to substances from the diet and the commensal microbiota, as well as potentially noxious compounds and pathogens. Coordination of productive host immune responses is a complex process that involves the integration of a disparate array of signals, not only from the pathogen but from the host as well. The gastrointestinal tract is highly innervated by intrinsic and extrinsic neurons to regulate not only physiology, but also these immune processes. External innervation to the intestine, including the sympathetic neurons and sensory neurons that arise from the sympathetic chain ganglia and dorsal root ganglia of the spinal cord respectively, are well appreciated to interface with the ENS and other cells to affect physiological processes^9^. While sensory neurons expressing the polymodal nociceptor TRPPV1 aid the development of host responses to enteric bacterial infection^5,8^, the role of colonic sympathetic innervation during the T-cell responses to *C. rodentium* infection was unknown.

In support of a role for central nervous system neurons in coordinating the immune response during enteric bacterial infection, we identified activation of several brain regions during the peak of *C. rodentium* infection. This included the nucleus tractus solitarius (NTS) within the brainstem, a sensory relay receiving input from visceral organs through vagal afferents. These data are in keeping with prior studies that demonstrated increased neuronal activation in vagal ganglia during *C. rodentium* and *Campylobacter jejuni* infection^25,26^. As an integrative center that can induce physiological responses depending on information from the viscera, it is not surprising that both excitatory and inhibitory neurons project from the NTS to other brain regions including the hypothalamus and medulla^27,28^.

Our studies uniquely identified neurons within the rostral ventrolateral medulla (RVLM), a major sympathetic control center in the brainstem ^29–31^, that were activated during the peak of *C. rodentium* infection. Although neurons within this brain region were originally identified to regulate heart rate and blood vessel contractility^29^, peripheral sympathetic innervation and neurotransmitters can inhibit systemic inflammation during a variety of insults including models of sepsis^32–34^.

Here we identified a host-protective role for sympathetic neurons, with peripheral sympathectomy significantly increasing bacterial burden during *C. rodentium* infection. These effects were due to significantly attenuated expression of *Ifnγ* in colonic T-cells. Although it was previously reported that chemical sympathectomy increased resistance to *Listeria monocytogenes* (*L. monocytogenes*), the purported mechanism is unclear. Evidence for this effect is limited to increased IFNγ and TNFα production in stimulated splenic T-cells from non-infected intact and sympathectomized mice^34^. In addition, while reduced *L. monocytogenes* was observed in sympathectomized IFNγ or TNFα deficient mice^34^, it is unclear if this is simply due to increased burden in those cytokine knockout mice. Our findings of reduced *Tnfa* and *Il1b* mRNA expression in the colon of *C. rodentium*-infected sympathectomized mice suggest that select aspects of mucosal and systemic immune responses are uniquely affected by sympathetic innervation of these tissues. Characteristic of the host responses to enteric *C. rodentium* is the infiltration of Th1-polarized CD4+ T-cells. These cells that produce IFNγ are thus critical during *C. rodentium* infection as this cytokine regulates many gene targets for host defense and bacterial clearance. The reduced colonic *Ifnγ* expression and IFNγ-producing colonic CD4+ T-cells is not due to an inability of naïve T-cells to undergo Th1 differentiation, as no difference in IFNγ production was observed in *in vitro* MLN T-cells enriched from control or sympathectomized mice. Reduced colonic IFNγ in sympathectomized infected mice is correlated with decreased *Nos2* and *Nox1* expression, which could impede antimicrobial reactive oxygen species production and increase bacterial burden^35^. Reduced colonic *Ifnγ* in the sympathectomized infected mice was insufficient to increase pathophysiological changes to the colon, exemplified by comparable crypt hyperplasia between the intact and sympathectomized infected groups. These effects on the intestinal epithelia are well appreciated to occur in response to infection-induced inflammation and IFNγ from responding CD4+ T-cells^20^. Thus, the lack of an increased hyperproliferative state in infected sympathectomized mice suggests that the increased bacterial burden was able to induce other factors capable of increased IEC proliferation and crypt hyperplasia.

Suggestive of differential regulation of systemic immune responses by sympathetic neurons, selective surgical ablation of the splanchnic nerve significantly increased TNFα, IL-6, and IFNγ production in rodent models of endotoxemia^36,37^ and reduced the abundance of live bacteria in the blood of sheep challenged with *E. coli*^34^. Further highlighting these negative regulatory effects of sympathetic innervation on innate immune responses, experimentally induced peritonitis increased CCL2 (MCP-1) chemokine expression, and consequently monocyte recruitment occurred in sympathectomized mice^38^.

Although the reasons for these differences in the response to systemic bacteremia and a non-invasive enteric bacterial pathogen are unknown, divergent responses could be simply due to differences in adrenergic receptors activated. Inhibition of inflammation by sympathetic innervation or neurotransmitters such as norepinephrine has been shown to require the β_2_AR subtype. Indeed, activation of C1 neurons within the RVLM prevented immunopathology in a model of ischemia-reperfusion injury and required β_2_AR signaling to activate the cholinergic anti-inflammatory pathway (CAIP)^39^. This anti-inflammatory reflex requires a highly specialized subset of T-cells that express the enzyme choline acetyltransferase (ChAT) that releases acetylcholine in response to β_2_AR stimulation^1,40,41^. Despite this clear role in reducing inflammation during systemic inflammation, ChAT+ T-cells aid the mucosal immune response during *C. rodentium* infection^17^. These prior studies highlight the diverse functions of neuroimmune circuits and the constituent cell types, which can be dependent on the threat and location of the response. In the colon, activation of local sympathetic neurons reduced the severity of induced colitis by decreasing expression of mucosal addressin cell-adhesion molecule-1 (MAdCAM1) ^42^. Although reduced MAdCAM1-mediated colonic infiltration of immune cells was suggested as the mechanism of action, no specific immune cell population or adrenergic receptor type was uniquely identified^42^.

Sympathetic inhibition of inflammation that occurs in models of sepsis has also been elucidated to be dependent not on the CAIP, but on the induction of IL-10 expression^32,36^. In the context of systemic infection, blockade of this pathway not only reduces IL-10 production but bacteremia as well^34^. The role of sympathetic innervation during infection is likely to depend on the nature of the pathogen. In support of this idea, enteric infection with *S. Typhimurium* activates sympathetic neurons that are found in close proximity to muscularis macrophages^20^. This neuron-macrophage communication was facilitated by β_2_AR signaling, which was proposed to induce an alternatively activated macrophage program and a tissue-reparative functionality^20^. Local sympathetic innervation of other mucosal surfaces, including the lung, has also been demonstrated to reduce innate inflammation induced by TLR agonists^43^. This innervation purportedly reduced inflammation and innate immune cell recruitment in a β_2_AR signaling- dependent manner, although the precise mechanism was not described^43^. With our data showing that βAR pharmacological antagonism, or β_2_AR KO mice have increased bacterial burden without reducing *Ifn*γ expression, the immune function of sympathetic neurotransmitters is highly dependent on the nature of the inflammatory or infectious agent.

Despite these immunosuppressive effects, proinflammatory roles of sympathetic neurons and neurotransmitters released have also been described. These contradictory roles have been attributed to the activation of αAR on target immune cells^44^. Activation of α_2_AR has long been appreciated to augment LPS-induced TNF production from macrophages *in vitro*^45^. Confirming this role of α_2_AR in enhancing inflammation, antagonism of this receptor subtype attenuated TNFα production during induced endotoxemia^46,47^. Adding further evidence of this proinflammatory role, intestinal inflammation evoked by dextran sulphate sodium was significantly attenuated in α_2_AR KO mice, and treatment of WT mice with α_2_AR agonists aggravated induced colitis^22^. From a pharmacology perspective, αAR activation by locally released norepinephrine from sympathetic neurons is logical, as norepinephrine has a significantly higher affinity for αAR compared to β_2_AR. Although the expression of αAR subtypes on immune cells has been previously described, the expression of αAR on CD4+ T-cells and the contribution to host responses during enteric bacterial infection were unknown. During *L. monocytogenes* infection, treatment with phentolamine mesylate, a non-selective αAR antagonist, significantly increased pathogen load in the liver, along with TNFα, IL6, and IFNγ production^48^. Our observations suggest that optimal T-cell responses require peripheral sympathetic neurons and αAR during enteric bacterial infection. Pharmacological antagonism of αAR significantly increased fecal and colonic *C. rodentium* and decreased colonic *Ifnγ* expression. Highlighting the unique effects of αAR signaling during infection, no effect on *Ifnγ* expression was observed with pharmacological blockade of βAR. These pharmacology studies using non-selective βAR antagonists were further supported by the lack of effect on infection-induced IFNγ production in β_2_AR KO mice. Together, these findings demonstrate the host immune response is uniquely controlled through sympathetic innervation and αAR signaling. To determine which populations of immune cells could be affected by this signaling axis, FACS-sorted cells from the colonic lamina propria were assessed for mRNA expression of known αAR subtypes. Expression of αAR subtypes on T-cells are typically low under baseline conditions and could be induced during T-cell activation^49^. Consistent with these data, we found that sorted colonic T-cells have significantly elevated expression of various αAR subtypes during *C. rodentium* infection, accompanied by dramatically increased IFNγ and IL17A production indicative of Th1 and Th17 responses respectively. Although increased T-cell proliferation has been reported upon αAR blockade^50,51^, colonic T-cell numbers were comparable between the non-infected intact and sympathectomized mice, implying that αAR signaling could be crucial for mediating host immune responses by altering T-cell function. Our data further indicates that although neutrophils, stromal cells, and LEC express these αAR subtypes, infection does not further increase the expression of these receptors. In the stromal cell compartment, only BEC expression of select αAR subtypes were found to increase with *C. rodentium* infection. To be clear, these data do not indicate a lack of biologically relevant function of the other receptor types despite failing to increase expression during infection.

Together our data demonstrate that peripheral sympathetic innervation and αAR are crucial for coordinating T-cell responses during *C. rodentium* infection. Control of mucosal immune responses in this manner further appears to be unique compared to the previously established effects on the systemic immune response. Finally, deciphering the role of αAR subtypes on mucosal immune cells during enteric bacterial pathogen infection may not only highlight novel approaches to enhance host protection but potentially limit immunopathology.

## Abbreviations

RVLM: rostral ventrolateral medulla
ENS: enteric nervous system
C. rodentium: *Citrobacter rodentium*
IEC: intestinal epithelial cells
ILC: innate lymphoid cells
Nos2: nitric oxide synthase
βAR: β-adrenergic receptors
αΑR: α-adrenergic receptors
β_2_AR KO: β_2_- adrenergic receptors knockout
LB: Luria Broth
CFU: colony-forming units
4-OHT: 4-hydroxytamoxifen
6OHDA: 6-hydroxydopamine
NTS: nucleus tractus solitarius
TH: Tyrosine Hydroxylase
d.p.i.: days post-infection
IEC: intestinal epithelial cells
CAIP: cholinergic anti-inflammatory pathway
ChAT: choline acetyltransferase
MLN: mesenteric lymph node
IFNγ: Interferon-γ
IL-17: interleukin-17
IL-22: interleukin-22

