## Supplemental figures for "The sympathetic nervous system enhances host immune responses to enteric bacterial pathogens"

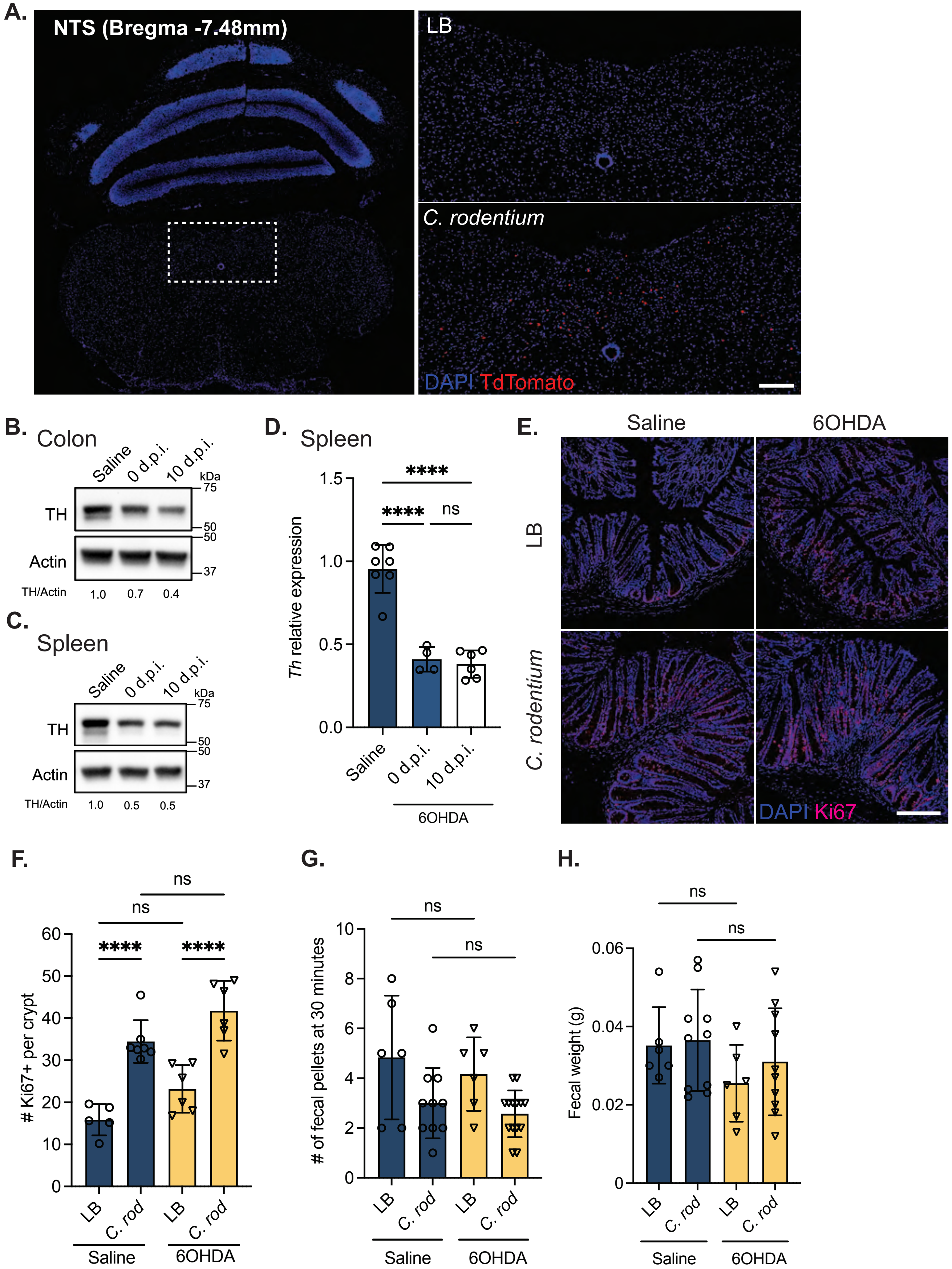


**Figure S1. *C. rodentium* infection activates discrete brain regions and histopathology is not altered by sympathectomy.** Representative images of the nucleus tractus solitarius (NTS, Bregma -7.48 mm) in LB and *C. rodentium* infected *Arc*TRAP mice at 10 days post-infection (d.p.i.) with activated neurons indicated by DAPI+ tdTomato+ cells **(A)**. Validation of sympathectomy by western blot showing expression Tyrosine Hydroxylase (TH) protein in the colon **(B)** and spleen **(C)**, or *Th* mRNA in the spleen **(D)** of mice treated with saline or 6OHDA 0 and 10 days post-infection (d.p.i.). Confocal microscopy conducted on colonic tissue sections for proliferating (DAPI+ Ki67+) intestinal epithelial cells **(E)** and the quantification of these cells **(F)** at 10 d.p.i. Distal colonic motility measured by quantifying the number of fecal pellets produced within a 15-minute period **(G)** and the weight of fecal pellets **(H)** in vehicle or 6OHDA-treated uninfected or infected mice. Results are from individual mice, mean ± SD. ns = not significant, **** P ≤0.0001. One-way ANOVA with Tukey's post-hoc test. Scale bar = 100 μm.

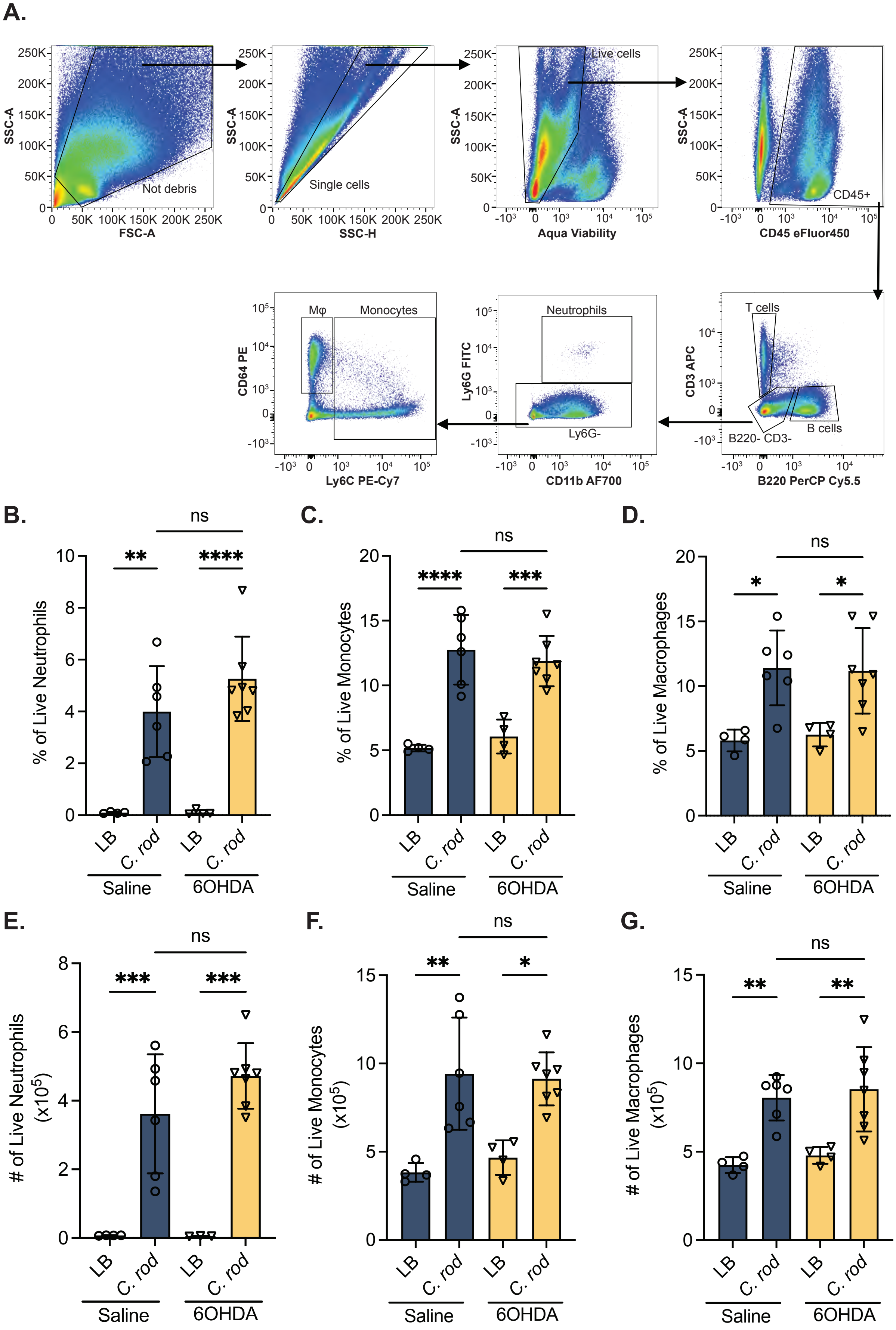


**Figure S2. Characterization of immune response in the sympathectomized *C. rodentium* infected mice.**
Flow cytometry was used to assess the immune responses during infection with the indicated gating strategy for the following immune cell populations: CD3+ T-cells, B220+ B-cells, Ly6G+ CD11b+ Neutrophils, Ly6C- CD64+ macrophages and Ly6C+ CD64- monocytes **(A)**. Using this approach, the frequency of live and number of neutrophils **(B&E)**, monocytes **(C&F)**, and macrophages **(D&G)** in the lamina propria of the colon of saline or 6OHDA-treated mice at 10 days post-infection (d.p.i.). Results are from individual mice, mean ± SD. ns = not significant, * P≤0.05, ** P ≤0.01, *** P ≤0.001, **** P ≤0.0001. One-way ANOVA with Tukey's post-hoc test.

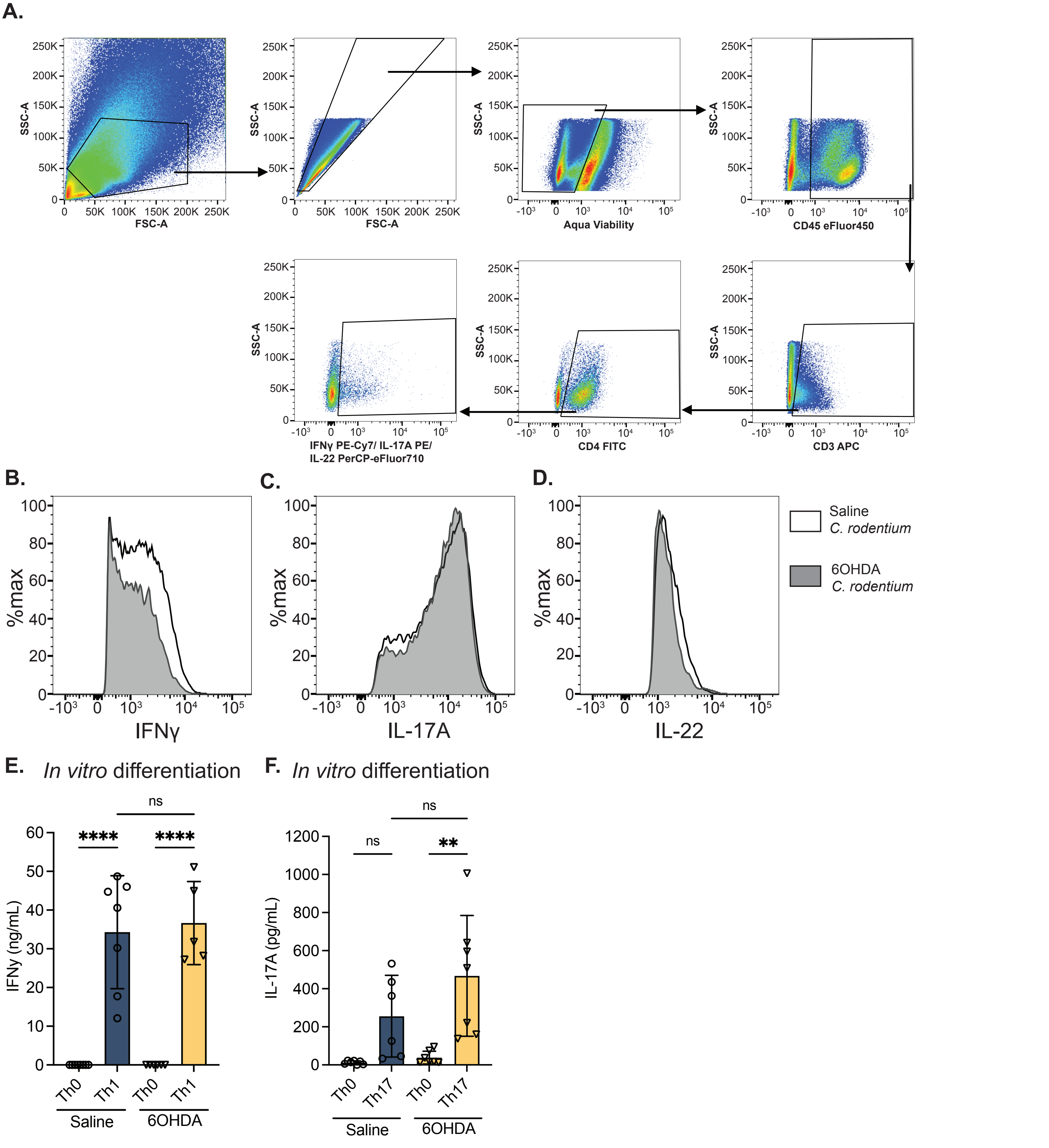


**Figure S3. Analysis of T-cells during *C. rodentium* infection in the control or sympathectomized mice.**
Flow cytometry was used with the gating strategy indicated to assess IFNγ, IL-17A, and IL-22-producing T-cells in the lamina propria of the colon **(A)**. Histograms showing intracellular cytokine staining for colonic CD3+ CD4+ T-cells that express IFNγ **(B)**, IL-17A **(C)**, and IL-22 **(D)** by flow cytometry in intact (white) and sympathectomized (grey) mice. Isolated CD4+ T-cells from the mesenteric lymph node of saline and 6OHDA treated mice were cultured in vitro with or without Th1 or Th17 differentiation media, and assessed for IFNγ **(E)** and IL-17A **(F)** production by ELISA. Results are from individual mice, mean ± SD. ns = not significant, * P≤0.05, ** P ≤0.01. One-way ANOVA with Tukey's post-hoc test.

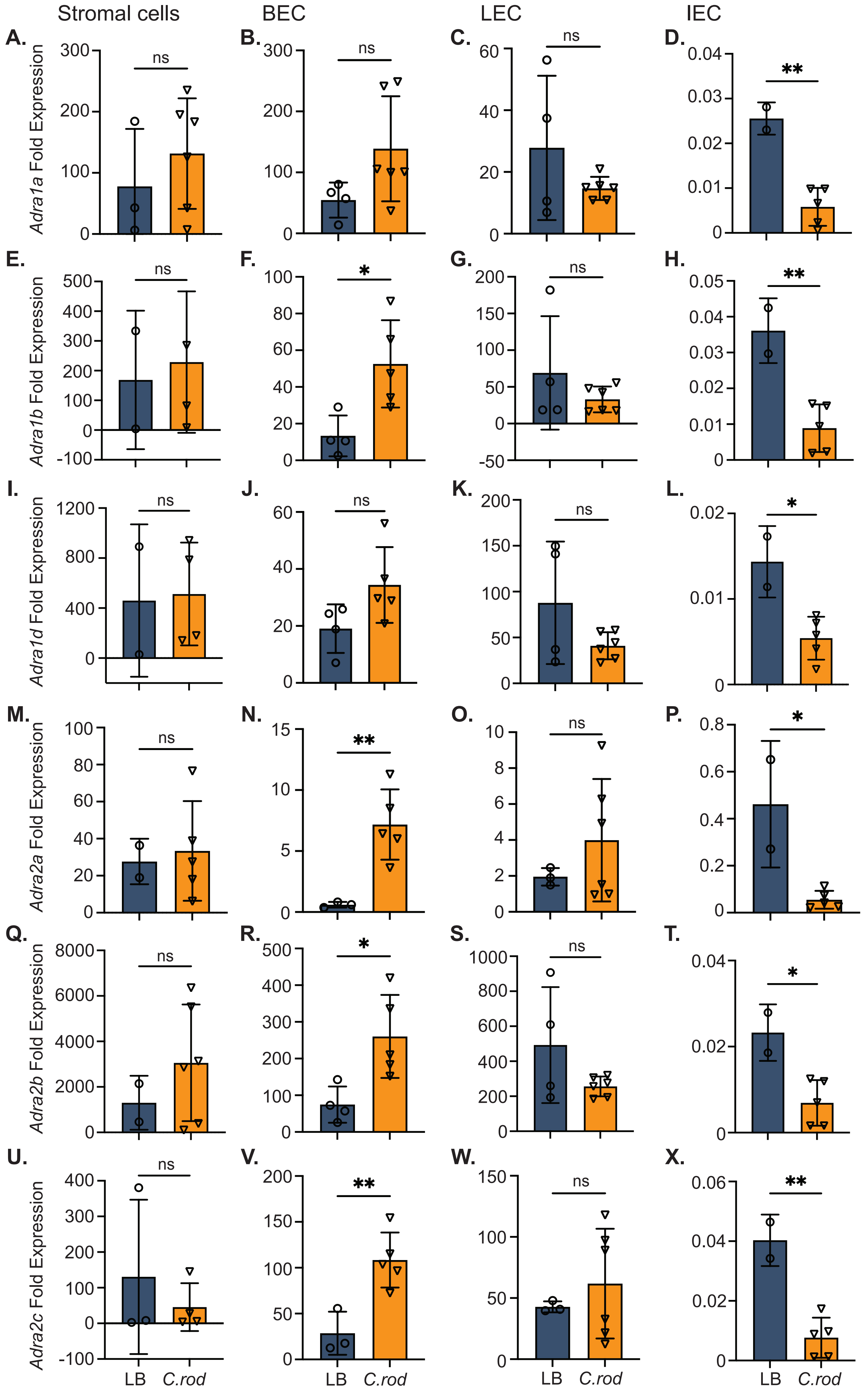

Stromal cells
BEC
LEC
IEC
A.
B.
C.
D.
E.
F.
G.
H.
I.
J.
K.
L.
M.
N.
O.
P.
Q.
R.
S.
T.
U.
V.
W.
X.
Adra1a Fold Expression
Adra1b Fold Expression
Adra1d Fold Expression
Adra2a Fold Expression
Adra2b Fold Expression
Adra2c Fold Expression
ns
*
**
LB
C.rod

**Figure S4. Expression of α-adrenergic receptors in colonic stromal and intestinal epithelial cells.**
The expression of α-adrenergic receptors encoded by *Adra1a* **(A-D)**, *Adra1b* **(E-H)**, *Adra1d* **(I-L)**, *Adra2a* **(M-P)**, *Adra2b* **(Q-T)** and *Adra2c* **(U-X)** was assessed in FACS-sorted colonic stromal cells (live, CD45-, CD31-), blood endothelial cells (live, CD45- CD31+ gp38-, 'BEC'), lymphatic endothelial cells (live, CD45-, CD31+ gp38+, 'LEC') and intestinal epithelial cells (IEC) from LB and *C. rodentium* infected mice at 10 days post-infection. Results are from individual mice, mean ± SD. ns = not significant, * P≤0.05, ** P ≤0.01. One-way ANOVA with Tukey's post-hoc test.

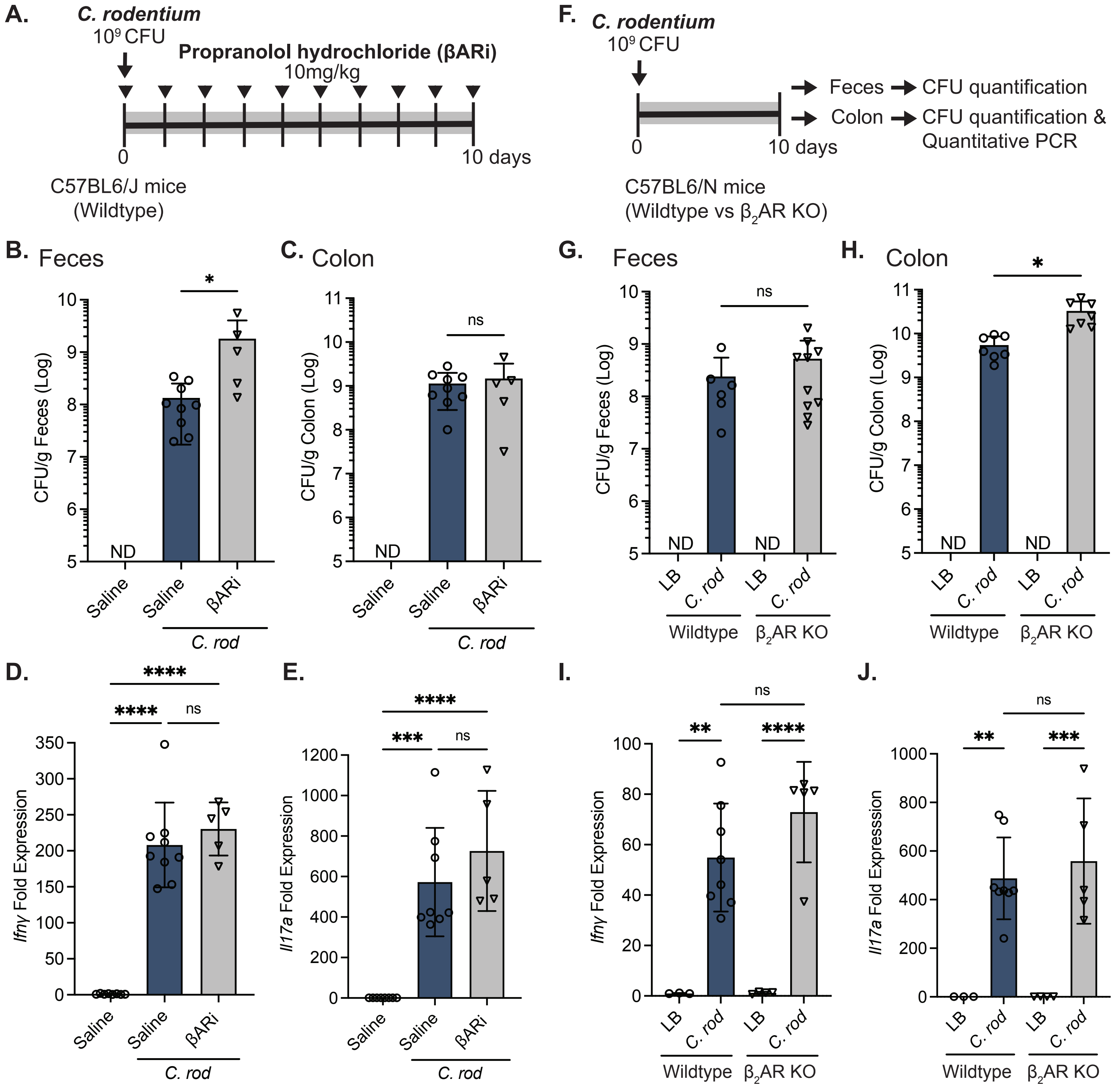


**Figure S5. Host IFNγ responses to *C. rodentium* in the colon do not require β-adrenergic receptor signaling.**
To assess the role of β adrenergic receptor (βAR) signaling during *C. rodentium* infection, the non-selective βAR antagonist (β ARi), Propranolol hydrochloride (10 mg/kg, i.p. one dose daily), was administered after *C. rodentium* infection and tissues collected 10 days post-infection (d.p.i.) **(A)**. Bacterial burden was determined by quantification of colony-forming units from feces **(B)** and colon **(C)**. Colonic mRNA expression of *Ifnγ* **(D)** and *Il17a* **(E)** in uninfected and infected saline or βARi-treated mice at 10 d.p.i. were assessed by RT-qPCR. The role of β2AR signaling was assessed by using wildtype and β2AR knockout (KO) C57BL6/N mice that were administered LB or C. rodentium followed by tissue collection 10 d.p.i. Quantification of colony-forming units of C. rodentium in feces **(G)** and colon **(H)**, as well as the colonic mRNA expression of *Ifnγ* **(I)** and *Il17a* **(J)** in uninfected and infected, wildtype and β2 AR KO mice at 10 d.p.i. Results are from individual mice, mean ± SD. ns = not significant, * P≤0.05, ** P ≤0.01, *** P ≤0.001, **** P ≤0.0001. One-way ANOVA with Tukey's post-hoc test.
